# Substrate recognition by the human mitochondrial processing peptidase and its processing of PINK1

**DOI:** 10.1101/2025.03.24.645073

**Authors:** Andrew N. Bayne, Danielle M. Simons, Naoto Soya, Gergely L. Lukacs, Jean-François Trempe

**Affiliations:** Department of Pharmacology and Therapeutics, McGill University, Montréal, QC H3G 1Y6, Canada; Centre de Recherche en Biologie Structurale, McGill University, Montréal, QC H3G 0B1, Canada; Department of Physiology, McGill University, Montréal, QC, Canada; Department of Biochemistry, McGill University, Montreal, Quebec, Canada

## Abstract

Nuclear-encoded mitochondrial proteins rely on N-terminal targeting sequences (N-MTS) for their import. These N-MTSs interact with the translocation machinery and are cleaved in the matrix by the mitochondrial processing peptidase (MPP), a heterodimeric zinc metalloprotease which is essential for the maturation of imported proteins. Import and cleavage of PINK1, a kinase implicated in Parkinson’s disease, govern its ability to sense mitochondrial damage, but the MPP cleavage site and its role in PINK1’s function remains cryptic. MPP typically cleaves a unique motif in N-MTSs with an arginine in the P2 position, but how MPP recognizes this motif is unclear. Here, we show that recombinant human MPP cleaves PINK1 between Ala28 and Tyr29 yet is turned over slowly compared to other canonical N-MTSs. While MPP cleavage is not required for downstream PARL processing or PINK1 accumulation in cells, the PINK1 N-MTS binds potently to MPP and inhibits the cleavage of other N-MTSs by glueing the regulatory (α) and catalytic (β) subunits. Finally, we utilize hydrogen-deuterium exchange mass spectrometry to reveal the binding site of the PINK1 N-MTS on MPP. Taken together, our work provides key insight into both the PINK1 import pathway and the mechanisms of MPP processing.

## Introduction

Mitochondria are multifaceted signalling hubs that regulate an array of cellular processes from oxidative phosphorylation to immunity, proteostasis, and more (Baker *et al*, 2011; Collier *et al*, 2023; Mills *et al*, 2017). While mitochondria contain their own genome, 99% of resident mitochondrial proteins are encoded in the nucleus and must be translated and imported into mitochondria from the cytosol (Busch *et al*, 2023). As such, mitochondria rely on a diverse set of translocases, chaperones, and proteolytic machinery to coordinate the import and processing of these imported substrates. The most common class of imported proteins feature an N-terminal mitochondrial targeting sequence (N-MTS), which tend to form amphipathic helices (Roise *et al*, 1988; Vögtle *et al*, 2009). These N-MTS proteins are guided via chaperones to the outer membrane of mitochondria (OMM), where they interact with the translocase of the outer membrane (TOM) complex (Mihara & Omura, 1996; Su *et al*, 2022). Upon exit from the TOM pore, N-MTS precursors interact with components of the translocase of the inner membrane 23 complex (TIM23) in the intermembrane space (IMS) and inner mitochondrial membrane (IMM) (Araiso *et al*, 2019; Callegari *et al*, 2020). At this stage, most N-MTS precursor proteins will either be sorted into the mitochondrial matrix, or laterally released into the IMM if they contain a hydrophobic stop-transfer signal (Mokranjac & Neupert, 2010; van der Laan *et al*, 2007). This substrate fate between IMM release or matrix localization hinges on the presequence-translocase associated import-motor (PAM) complex associated to TIM23, as well as the mitochondrial membrane potential (Schendzielorz *et al*, 2018). In both cases, the N-terminus of the imported protein can be matrix-exposed and accessible for cleavage.

In the matrix, these N-MTSs are cleaved off by the mitochondrial processing peptidase (MPP), a heterodimeric metalloprotease consisting of MPPα and MPPβ subunits (Gakh *et al*, 2002). While both subunits are required for MPP processing activity *in vitro* (Saavedra-Alanis et al, 1994), each monomer imparts a distinct role during MTS binding and cleavage: MPPβ coordinates Zn^2+^ via a reversed thermolysin binding motif (HXXEH) and is responsible for cleaving the scissile bond within the MTS (Luciano *et al*, 1998; Taylor *et al*, 2001); MPPα is a pseudoprotease that does not coordinate Zn^2+^ and instead mediates early MTS binding and recruitment, primarily via its glycine rich loop (GRL) (Dvořáková-Holá *et al*, 2010; Nagao *et al*, 2000; Shimokata *et al*, 1998). After MTS removal by MPP, precursor proteins may be pulled into the matrix to fold with the matrix chaperone machinery, or they may undergo subsequent proteolytic steps before reaching their mature form (Deshwal *et al*, 2020). This processing by MPP is essential for the folding and function of its mitochondrial substrates. Indeed, knockout of either *PMPCA* (encoding MPPα) or *PMPCB* (encoding MPPβ) in mice results in embryonic lethality (Blake *et al*, 2021). Mutations in *PMPCA* cause autosomal recessive ataxias (Choquet *et al*, 2016; Jobling *et al*, 2015; Takahashi *et al*, 2021), severe mitochondrial disease (Joshi *et al*, 2016), and encephalopathy (Rambani *et al*, 2023). Likewise, mutations in *PMPCB* lead to childhood neurodegeneration (Vögtle *et al*, 2018) and Leigh syndrome with ataxia (Matthews *et al*, 2024). MPP thus plays a critical, global role in maintaining mitochondrial health.

Our current understanding of MPP has been largely limited to studies of its yeast ortholog. The structure of yeast MPP was determined in 2002, revealing how MTS peptides bind to MPPβ in an extended conformation for proteolysis (Taylor *et al*., 2001). This study, along with others, confirmed the preferred recognition motif of yeast MPP: an arginine in the P2 position (*i.e.* N-terminal to the scissile bond) and a bulky, hydrophobic residue (denoted Φ) in the P1’ position (*i.e.* C-terminal to the scissile bond) (Ogishima *et al*, 1995). While these MPP cleavage motifs were also shown to be conserved in humans via broad mass spectrometry N-terminomics experiments (Calvo *et al*, 2017), the structural mechanisms of MTS binding to human MPP, particularly the substrate recognition role of MPPα, remain underexplored. This research gap also precludes our ability to study metazoan specific MPP substrates that have no yeast ortholog.

One such metazoan specific MPP substrate is PINK1 – a mitochondrial kinase whose mutations cause early-onset Parkinson’s disease (Valente, 2004). PINK1 functions as a sensor of mitochondrial damage by accumulating on depolarized mitochondria to initiate their selective engulfment by the autophagosome, a process coined mitophagy (Lazarou *et al*, 2015). Strikingly, mitochondrial import and proteolysis of PINK1 dictate its ability to sense mitochondrial damage: (1) full length PINK1 (64 kDa) localizes to the TOM complex via N-MTS, specifically via its amino acids 1-90, where it is threaded through the Tom40 pore (Okatsu *et al*, 2015). (2) The N-MTS of PINK1 is imported into the mitochondrial matrix via an atypical mechanism, resistant to Tim23 or PAM subunit mtHSP70 impairment (Filipuzzi *et al*, 2017; Sekine *et al*, 2019). (3) The N-MTS binds to and is cleaved by MPP at an unknown residue (Greene *et al*, 2012). (4) The transmembrane domain (TMD) of PINK1 is cleaved by PARL between Ala103 and Phe104, which generates the 52 kDa PINK1 fragment that is retrotranslocated to the cytosol and degraded by the proteasome (Deas *et al*, 2011; Jin *et al*, 2010; Yamano & Youle, 2013). In the absence of damage, PINK1 is thus turned over rapidly to prevent the errant removal of healthy mitochondria. Following mitochondrial stress, typically depolarization or misfolded protein accumulation in the matrix (Jin & Youle, 2013; Narendra *et al*, 2010), PINK1 accumulates as part of a TOM-TIM23 supercomplex (Akabane *et al*, 2023; Eldeeb *et al*, 2024), where it dimerizes and phosphorylates ubiquitin at Ser65 to generate phospho-ubiquitin (pUb) (Koyano *et al*, 2014; Okatsu *et al*, 2013; Rasool *et al*, 2022). In turn, pUb chains recruit the E3 ligase Parkin to start mitophagy (Sauvé *et al*, 2015).

Still, the determinants for PINK1 steady-state import remain cryptic, along with the roles of its N-MTS and MPP during this process. While the PARL cleavage site has been validated both in cells and using recombinant proteins *in vitro* (Lysyk *et al*, 2021), the cleavage site of the PINK1 N-MTS by MPP remains poorly defined aside from one study, which reveal an intermediate “MPP-cleaved” PINK1 fragment that appears on immunoblots upon knockdown of PARL (Greene *et al*., 2012). Overall, there is a critical need to characterize the human MPP dimer, both to shed light on the mechanisms of mitochondrial import and processing in mammals and to clarify the role of MPP in PINK1 function. In this work, we: produce recombinant human MPPαβ; elucidate the MPP cleavage site on PINK1; compare MPP proteolysis of PINK1 to other MTSs; develop a broadly applicable quenched fluorescent peptide to probe human MPP activity; and unveil the dynamics of MPP upon binding to the PINK1 MTS, providing a model for substrate recognition by human MPP. Together, our work provides key insight into the function of human MPP and its role in PINK1 import.

## Results and Discussion

### Recombinant human MPP can be purified from *E. coli* in an active form

First, we sought to develop a platform for producing recombinant human MPPαβ. To provide structural context for protein engineering, we generated a model of the human MPPα-MPPβ dimer using AlphaFold3 (Abramson *et al*, 2024) **(Fig 1A)**. Based on a previous report that rat MPPα was largely insoluble when purified as a monomeric MBP-fusion protein (Saavedra-Alanis *et al*., 1994), we attempted to purify human and mouse GST-MPPα and human GST-MPPβ from *E. coli*. While GST-MPPβ yielded soluble protein following GST elution, both human and mouse GST-MPPα resulted in negligible recovery of soluble GST-fusion protein **(Fig 1A)**. To this end, we opted for a co-expression system in which both human genes are expressed simultaneously to enhance MPPα folding and solubility. However, full-length mature MPPα-MPPβ did not express in *E. coli* until we truncated the hydrophobic and proline-rich N-terminus of MPPα **(Fig S1B)**, which is conserved across vertebrates, but not invertebrates or yeast **(Fig S2A)**. Still, small scale purifications of Δ60-MPPα-MPPβHis_6_ resulted in significant loss of untagged MPPα **(Fig S1B)**. We therefore tagged the less soluble Δ60-MPPα subunit to maximize dimer recovery, which yielded reproducible MPPαHis_6_-MPPβ purifications upon scaling up, despite inclusion body formation **(Fig S1C)**. Gel filtration of affinity-purified Δ60-MPPα_His6_-MPPβ resulted in resolvable peaks corresponding to MPPαβ dimer and excess MPPα monomer, as visualized on SDS-PAGE **(Fig 1B)**, corroborating that our N-terminal deletion did not inhibit MPP dimerization. We also confirmed the masses of our recombinant human MPP using mass spectrometry and verified the purity of both MPPα_his6_β and MPPα_his6_ to be >90% **(Fig S3)**. Our yields of the human MPP dimer (30-50 μg/L of *E. coli* culture) were reproducible and enabled us to proceed with functional studies of human MPP and its processing of PINK1.

**Figure 1.**
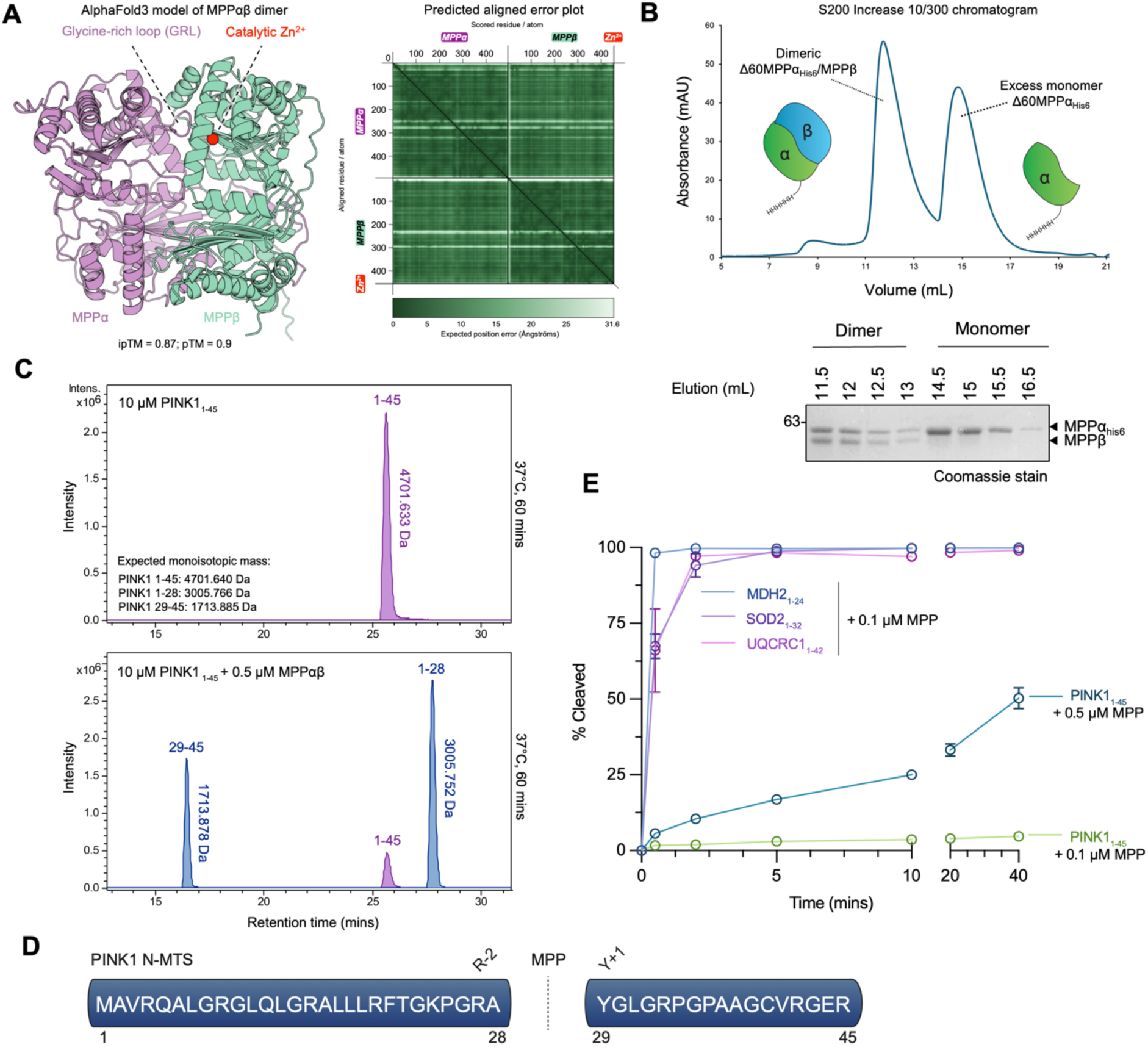
Recombinant MPPαβ cleaves PINK1_1-45_ between Ala28 and Tyr29 on a distinctly slow timescale. **A)** AlphaFold3 model of MPPαβ dimer (left), visualized in pyMOL and annotated with predicted template modeling (pTM) score and interface predicted template modeling (ipTM) scores. Predicted aligned error plot from AlphaFold3 visualized using PAE viewer (Elfmann & Stülke, 2023) (right). **B)** Size exclusion chromatogram (top) of Δ60-MPPα_his6_β resolving MPPαβ dimer from excess MPPα monomer. Fractions spanning both peaks were resolved on SDS-PAGE and visualized using Coomassie stain (bottom). **C)** LC-MS extracted ion chromatograms (EICs) of PINK1_1-45_ (purple) and its cleavage products (blue) following incubation with recombinant MPPαβ dimer at 37°C for 60 mins, annotated with observed and expected masses for each peptide. **D)** Schematic of PINK1_1-45_ and its MPP cleavage site between Ala28/Tyr29, yielding PINK1_1-28_ and PINK1_29-45_ fragments. **E)** Time course of MTS processing using synthetic peptide MTSs and recombinant MPPαβ as monitored by LC/MS. EICs were generated for each full-length MTS and their cleavage products. EIC intensities were exported and quantified as % cleaved. All data points were performed in duplicate (n=2) and were plotted as the mean % cleaved ± SD.

### MPP cleaves PINK1_1-45_ between Ala28 and Tyr29 on a distinctly slow timescale

To determine the MPP cleavage site on PINK1, we incubated MPPαβ with a synthetic peptide corresponding to the PINK1 N-MTS (a.a. 1-45) and monitored the reaction products using mass spectrometry. Extended incubation of MPP with PINK1_1-45_ led to the selective processing of PINK1 between Ala28 and Tyr29 **(Fig 1C)**, with Arg27 acting as the canonical “R-2” MPP cleavage motif **(Fig 1D)**. This is consistent with the migration shifts observed following PARL knockdown, which locates the cleavage within the first 40 a.a. (Greene *et al*., 2012). Next, we compared the kinetics of PINK1 processing to other canonical MTSs by monitoring MPP cleavage of various peptide substrates across time **(Fig 1E)**. In this assay, MDH2_1-24_, SOD2_1-32_, and UQCRC1_1-42_ were all 95%+ processed within 2 minutes after incubation with 0.1 µM MPP at 37 °C **(Fig 1E)**. Strikingly, only 5% of PINK1_1-45_ was cleaved under these conditions after 40 minutes. Upon incubation with 0.5 µM MPP, PINK1_1-45_ displayed significant processing by MPP, but was still slower than the other substrates and was only 50% cleaved after 40 minutes **(Fig 1E)**. While it was previously shown in yeast MPP that longer MTSs are more dependent on MPPα for their processing (Kitada *et al*, 2003), the slow turnover of PINK1_1-45_ cannot be explained only by the length of its MTS, given that UQCRC1_1-42_ was still rapidly turned over. This difference points to the presence of unique determinants within the PINK1 MTS that alter its binding to and processing by MPP.

Since PINK1 harbors high MTS propensity across its first 95 amino acids (Bayne *et al*, 2023), which also contains multiple possible R-2/Φ+1 motifs (namely R88/W91), we asked whether there may be a second upstream cleavage site that would be processed on a more rapid timescale, like other canonical MTSs. We also interrogated the effect of mutating R27, given that previous studies in yeast had shown that mutation of R-2 motifs should severely reduce MTS cleavage by MPP (Niidome *et al*, 1994). To this end, we performed a cleavage assay comparing PINK1_1-45_, PINK1_1-45(R27A)_, PINK1_1-35_, PINK1_35-60_, and PINK1_61-95_ (three peptides which span the entirety of the PINK1 N-terminus) **(Fig 2A)**. In this assay, both PINK1_1-45_ and PINK1_1-35_ were reliably processed by MPP between A28/Y29, while mutation of R27 abrogated MPP cleavage, as predicted. Both PINK1_35-60_ and PINK1_61-95_ did not display any significant cleavage by MPP, effectively confirming the singular MPP cleavage site on PINK1 within the first 35 amino acids.

**Figure 2.**
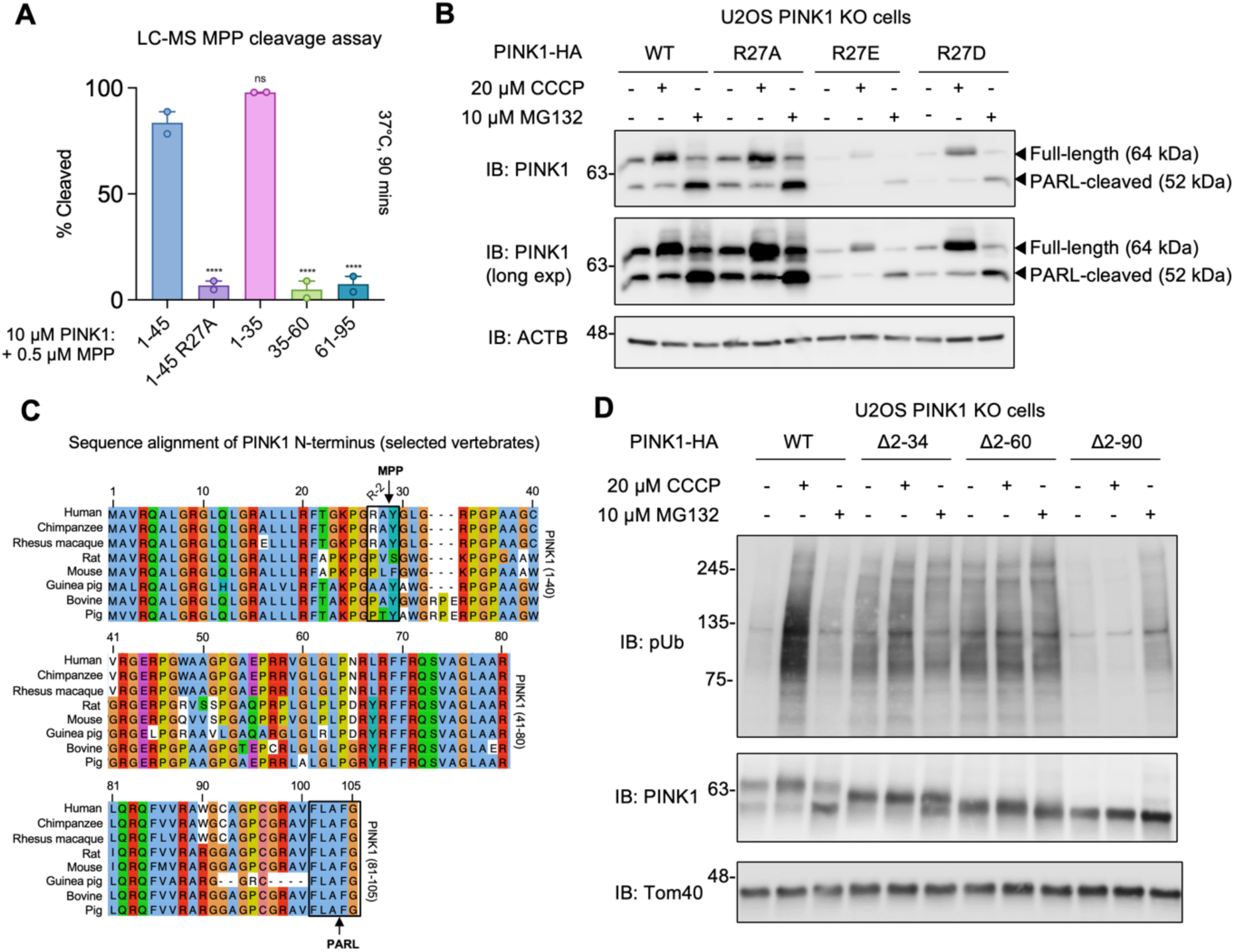
MPP uniquely cleaves the PINK1 N-terminus at Ala28/Tyr29 but is not a prerequisite for PARL proteolysis in cells. **A)** Comparison of processing for various PINK1 MTS peptides after 90 min incubation with MPP at 37°C. EICs were generated and processed as in Figure 1E. Data points represent the mean % cleaved ± SD (n=2). One-way ANOVA followed by Dunnett’s test were performed to compare the % cleaved of each reaction to PINK1_1-45_ as control (ns = P > 0.05, * = P ≤ 0.05, ** = P ≤ 0.01, *** = P ≤ 0.001, **** = P ≤ 0.0001). **B)** U2OS PINK1 KO cells were transfected with WT PINK1 or denoted mutants and treated with DMSO, 20 µM CCCP, or 10 µM MG132 for 3 hours. Cells were collected, lysed, normalized using the BCA assay, and subjected to immunoblotting with the indicated antibodies. **C)** Sequences of vertebrate PINK1_1-110_ were aligned using the MUSCLE algorithm in Jalview 2.11.4.1. **D)** U2OS PINK1 KO cells were transfected, treated with DMSO, CCCP at the indicated concentrations, or 10 µM MG132 for 3 hours and processed as in Fig 2B.

### MPP proteolysis of PINK1 is not required for downstream PARL cleavage

Given our findings of a singular, unique MPP cleavage site within the PINK1 N-terminus, we set out to resolve whether MPP cleavage of PINK1 is a pre-requisite for downstream PARL cleavage within its TMD. To this end, we transfected designer PINK1 mutants into U2OS PINK1 KO cells and treated them with carbonyl cyanide m-chlorophenyl hydrazone (CCCP) to depolarize mitochondria and stabilize full length PINK1 (64 kDa), or MG132 to stabilize PARL-cleaved PINK1 in the cytosol (52 kDa). First, we mutated the R-2 motif in PINK1, which flanks the scissile bond and mutation of which (*i.e.* R27A) resulted in the loss of MPP cleavage *in vitro* **(Fig 2A)**. PINK1 R27A was still cleaved by PARL in cells, as evidenced by the 52 kDa PINK1 band upon MG132 treatment **(Fig 2B)**. We also introduced charge reversals to the R-2 motif with R27E/R27D PINK1, which should further reduce any residual MPP processivity that R27A might harbor in cells. Still, PINK1 R27E/R27D behaved comparably to WT PINK1 by maintaining CCCP-responsiveness and generating the MG132-dependent, PARL-cleaved fragment **(Fig 2B).** These findings suggest that MPP proteolysis of PINK1 is not required for downstream PARL processing or for accumulating in response to CCCP-induced depolarization.

Nevertheless, PINK1 binding to MPP during import may still play a role in its ability to sense mitochondrial damage, independent of cleavage or R27 mutation. Consistent with this hypothesis, the PINK1 R-2 motif is poorly conserved across vertebrates, while the regions on either side of the scissile bond are highly conserved, as well as the PARL cleavage site **(Fig 2C)**. Deletion of PINK1 a.a. 2-34, as well as longer deletions 2-60 and 2-90, indeed abolish PINK1’s ability to respond to loss of membrane potential, instead accumulating constitutively on mitochondria (**Fig. 2D**), as observed previously (Okatsu *et al*., 2015). While basal accumulation of PINK1_Δ2-34_ may be driven by other import machinery, exploring MPP-PINK1 binding is necessary to provide additional context for its unique import behavior.

### PINK1_1-45_ competes potently with other substrates to inhibit MPP, as measured in IQF-FRET competition assays

To profile MPP processing against diverse substrates and to further characterize its interaction with PINK1, we established a FRET-based activity assay similar to an approach used to characterize yeast MPP (Ogishima *et al*., 1995). Briefly, internally quenched fluorescent (IQF) peptides with a cleavable sequence were synthesized with a fluorophore (2-aminobenzoyl (Abz)) and a quencher (*e.g.* 2,4-dinitrophenyl (Dnp)) positioned on either side of the scissile bond. When the substrate is intact, Abz fluorescence is quenched due to the proximity of the Dnp quencher. Cleavage increases the distance between the fluorophore-quencher pair, permitting Abz fluorescence emission **(Fig 3A)**. This enables the real-time monitoring of proteolytic reactions by measuring fluorescence emission (*i.e.* the cleaved Abz-product) over time.

**Figure 3.**
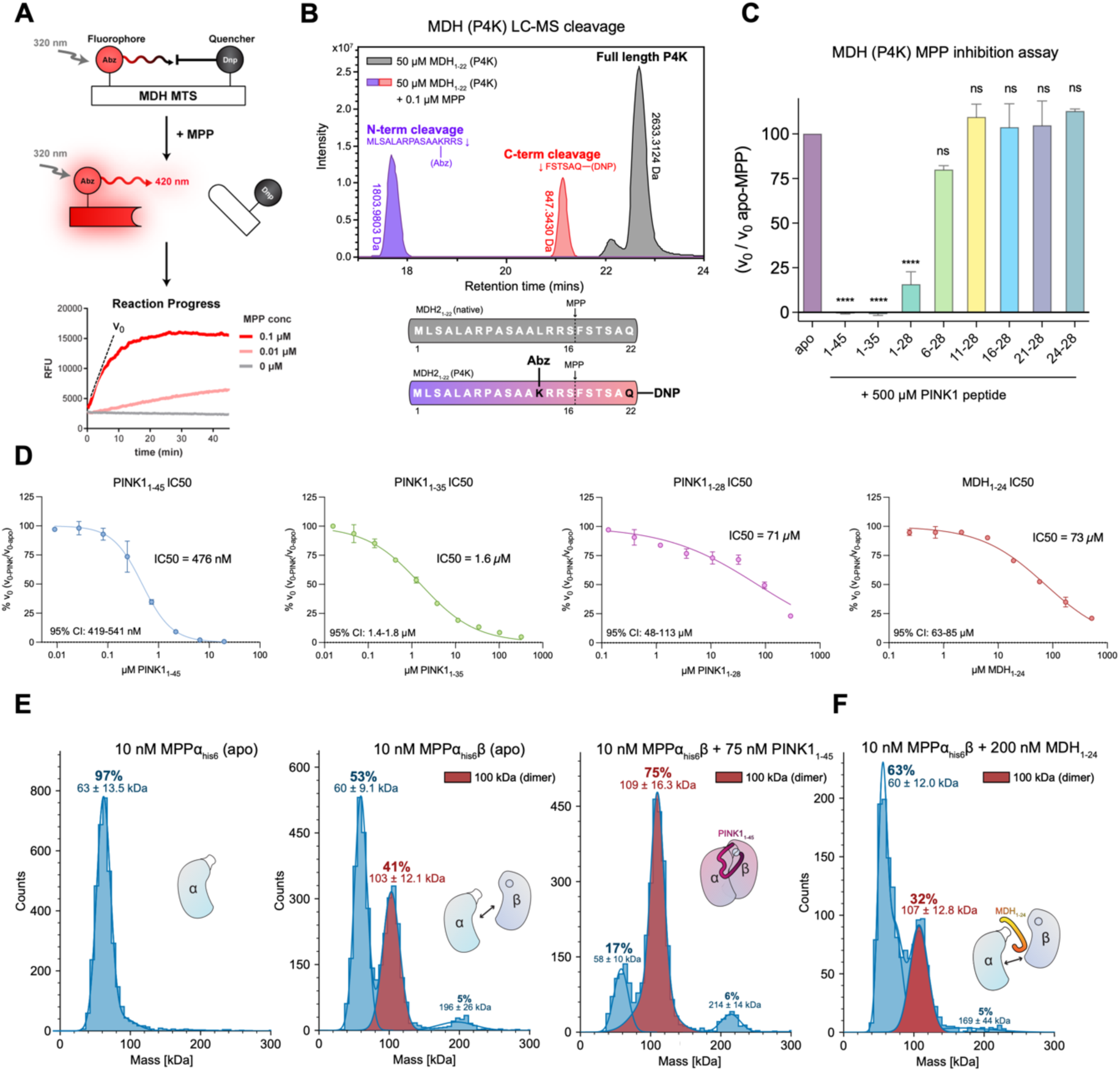
PINK1_1-45_ potently inhibits MPP processing and traps MPPαβ dimerization. **A)** Schematic of the workflow for the MDH2-based internally quenched fluorescent (IQF) peptide assay used to monitor MPP activity in Fig 2C and 2D. **B)** LC-MS characterization of P4K cleavage by MPP, visualized as in Fig 1C. **C)** 10 µM of P4K peptide ± 500 µM of indicated PINK1 peptides were incubated with 50 nM MPPαβ and reactions were monitored for 60 mins in a plate reader. Reaction velocities (v_0_) were calculated as the initial linear slope of each progress curve and were converted to % v_0_ by dividing each v_0_ by the mean of the apo-MPP v_0_. Data are plotted as the mean relative v_0_ ± SD (n=2). Statistical analysis was performed as in Fig 2A. **D)** MTS peptides were titrated into IQF-FRET assay reactions at various concentrations for IC_50_ determination. At each concentration, the relative inhibition was calculated as in Fig 2C. Non-linear IC_50_ curve-fitting was performed in GraphPad Prism 10.2.1. Data points represent the mean relative v_0_ ± SD (n=2). **E)** Monomer-dimer proportions of 10 nM MPPα or MPPαβ were measured on a Refeyn TwoMP mass photometer, either apo- or immediately following addition of PINK1_1-45_ peptide. Mass histograms were binned, fitted with Gaussian curves, and analyzed in DiscoverMP software (v2024 R1, Refeyn). **F)** Same as Fig 3E with MDH_1-24_.

To implement this approach, we designed two IQF peptides based on the human MDH2 (a.a. 1-22) MTS sequence: one with N-terminal Abz fluorophore and C-terminal Dnp quencher, and one with Leu13 (in the P4 position of the cleaved bond) mutated to Lys-Abz and C-terminal Dnp, referred to as “P4K” or “MDH-P4K” in this text **(Fig S4A)**. While both substrates were cleaved by MPP in a concentration-dependent manner, the reduced proximity between Abz/Dnp in the P4K substrate resulted in a higher dynamic range of fluorescence from baseline to complete proteolysis **(Fig S4A)**, and as such, we proceeded using the P4K peptide. Using mass spectrometry, we confirmed the canonical R-2 cleavage site within the MDH2-P4K peptide, between Ser17 and Thr18 **(Fig 3B)** and estimated the kinetic parameters of the P4K peptide to be K_M_ = 4.6 µM (95% CI: 2.3 to 9.1) and V_max_ = 0.058 µM/sec (95% CI: 0.048 to 0.069) **(Fig S4B)**. Using our P4K-FRET assay, we confirmed that human MPPαβ is only active when both subunits are present, that is, neither monomeric MPPα nor MPPβ were able to cleave MDH-P4K **(Fig S4C)**. As expected, human MPPαβ processing was also abrogated by addition of EDTA in a dose-dependent manner **(Fig S4D)**. We screened the effect of divalent cation addition on MPP activity, given a previous report that molar excess of Zn^2+^ inhibited yeast MPP processing with a K_i_ of 3.1 µM (Luciano *et al*., 1998). Remarkably, Zn^2+^ was also the only cation which significantly inhibited human MPP in our screen **(Fig S4C)**, yielding a congruent IC_50_ of 3.6 µM (95% CI: 2.3-5.5 µM) **(Fig S4D)**. Together, these results show that the IQF-FRET assay reliably captures MPP’s cleavage activity and could be exploited to screen for inhibitors or modulators.

Next, we screened how various truncations of the PINK1 N-MTS would inhibit the processing capacity of the P4K substrate. We find that 500 µM PINK1_1-45_ or PINK1_1-35_ resulted in a complete inhibition of P4K processing, as measured by the initial velocity (*v_0_*) compared to apo-MPP against P4K in the absence of any competing peptides **(Fig 3C)**. The cleavage product PINK1_1-28_ also inhibited MPP processing of P4K, albeit to a lesser degree. Removal of the first 5 N-terminal amino acids resulted in a significant decrease in the inhibitory capacity of the PINK1 N-MTS, suggesting that these residues are critical in anchoring PINK1 to MPP and inhibit cleavage. All other N-terminally truncated peptides did not inhibit MPP at concentrations of 500 µM. Following this broad screen, we used dose response curves to determine the potency for PINK1_1-45_, PINK1_1-35_, and PINK1_1-28_, yielding IC_50_’s of 476 nM, 1.6 µM, and 71 µM, respectively **(Fig 3D)**. In the same assay, MDH2_1-24_ displayed an IC_50_ of 73 µM, suggesting that PINK1_1-45_ binds more potently to MPP, and that this potency is driven by both proximal and distal portions of the PINK1 N-MTS, on either side of the scissile bond.

### PINK1_1-45_ acts as a molecular glue of active MPPαβ dimers

To further contextualize the potency and slow turnover of the PINK1_1-45_ peptide, we asked how PINK1_1-45_ binding might alter MPPαβ complex formation, either by preventing or augmenting MPPαβ dimerization. To answer this, we utilized mass photometry, a bioanalytical technique that measures the molecular mass of native complexes using single biomolecule interferometric scattering microscopy. In this assay, the gel filtration peak of monomeric MPPα_his6_ displayed a uniform distribution, with nearly 100% of the particles at 63 ± 13.5 kDa, corresponding to the expected mass of monomeric MPPα **(Fig 3E)**. The dimeric MPPαβ peak, which was resolved at 4 °C on gel filtration, revealed that ∼53% of MPPαβ was monomeric in the absence of substrate and only ∼41% MPPαβ was dimeric. Given that these mass photometry experiments were conducted at room temperature using 10 nM of each protein, this suggests that apo-MPP is in dynamic exchange between monomer and dimer, with a dimerization constant in the nM range. Upon addition of PINK1_1-45_ in 7.5-fold molar excess, MPPαβ dimerization was promoted, increasing the dimeric MPP populations to 75%. This enhancement of MPPαβ dimerization appears unique to the PINK1_1-45_ peptide, that is, MDH_1-24_ did not significantly alter the monomer-dimer distribution of active MPPαβ when added in 200-fold molar excess **(Fig 3F)**.

These findings posit PINK1 as a substrate trap or molecular glue of active MPPαβ and suggest that PINK1_1-45_ inhibits MPP by preventing the dissociation of MPP subunits (**Fig 3E**). MPPαβ may require dynamic interconversion between monomeric and dimeric states to facilitate multiple rounds of substrate binding and catalysis. Still, based on the potent binding **(Fig 3D)**, slow turnover **(Fig 1E)**, and dimer trapping capacity **(Fig 3E)** of PINK1_1-45_, we propose that PINK1_1-45_ serves as a model substrate to study time-resolved structural dynamics of the active MPPαβ complex and reveal the mechanisms that drive MPP substrate recognition and binding.

### Structural dynamics measured by HDX-MS reveal how MPPα orchestrates substrate recognition

To investigate the conformational rearrangements of MPP in response to substrate binding, we performed HDX-MS on MPPαβ in the absence and presence of the PINK1_1-45_ peptide and mapped the changes in deuterium exchange onto the AlphaFold structure of MPPαβ. To probe early stages of MPP-substrate binding, we incubated MPPα_his6_β with or without PINK1_1-45_ (6-fold molar excess) on ice for 1 min, followed by HDX deuterium labelling (10 sec, 1 min, and 5 min) and quenching. In this assay, we achieved nearly complete sequence coverage of both MPPα (98%) and MPPβ (99%) **(Fig S5A, S6A, Table S1)**. In general, the exchange rates after 5 min labeling agree with the structure predictions, with loops and termini displaying higher exchange rates. Strikingly, we observe that PINK1_1-45_ induces significant protection in several MPPα structural features **(Fig 4A, S5B)**, with less extensive differences observed in MPPβ in the vicinity of the active site **(Fig S6B)**.

**Fig 4.**
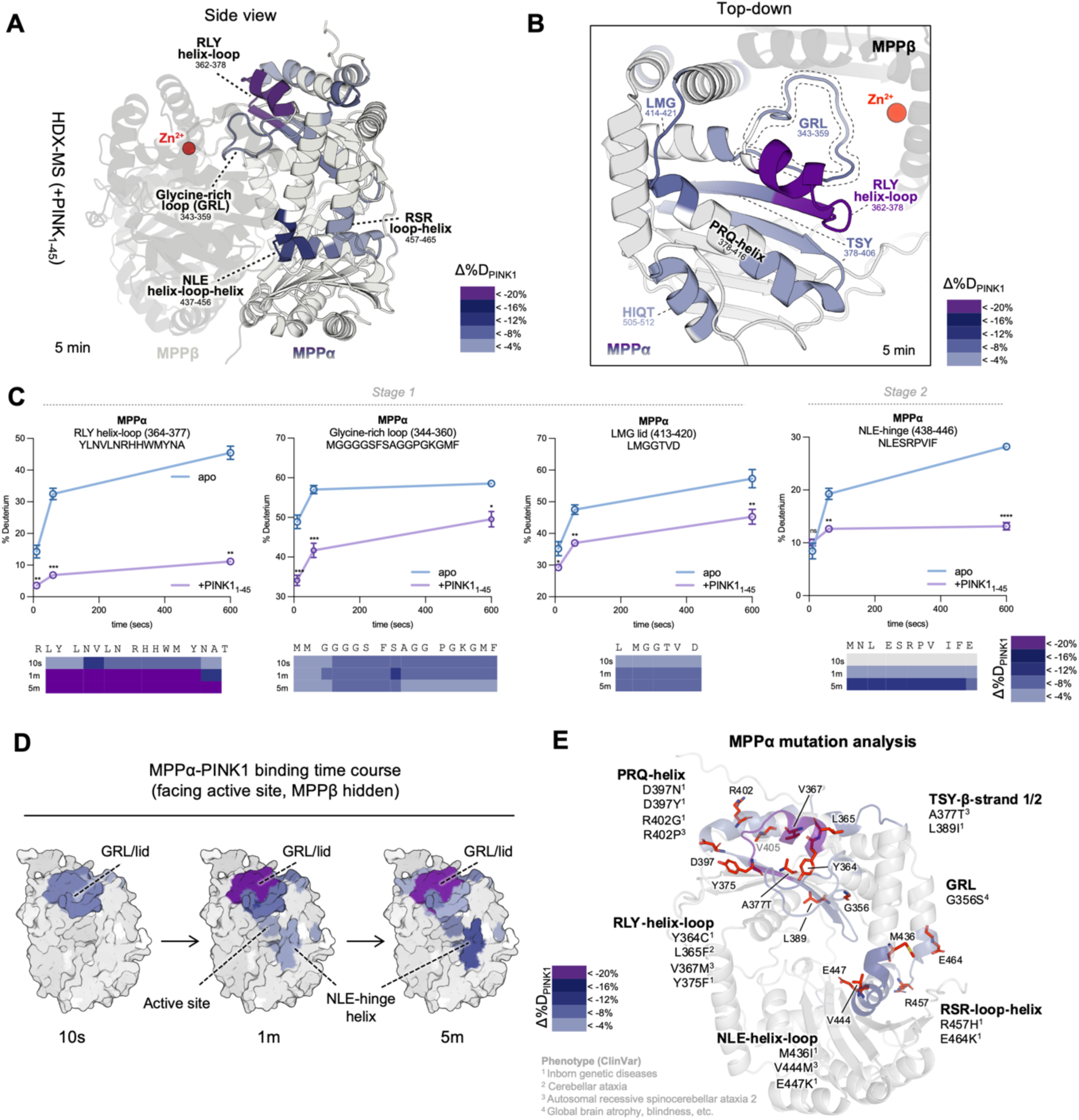
HDX-MS reveals the conformational dynamics of MPPα during PINK1_1-45_ binding. **A)** Structural mapping of per-residue deuterium exchange differences on MPPα after 5 minutes of D_2_O exposure ± PINK1_1-45_, visualized in PyMOL using the AlphaFold3 MPPαβ structure and coloured according to the Δ%D_PINK1_ scale bar. **B)** Top-down view of the MPPα lid, visualized as in Fig 4A. **C)** Deuterium uptake plots for selected regions of MPPα showing protection. Curves show percent deuterium incorporation in the absence (blue) and presence (purple) of PINK1_1-45_ across timepoints. All data points represent n=3 replicates ± SD. Statistical significance was determined using an unpaired two-tailed t-test with Welch’s correction (*P ≤ 0.05, **P ≤ 0.01, ***P ≤ 0.001, ****P ≤ 0.0001). Representative peptides are shown below each uptake plot, with coloured rectangles indicating % deuterium change according to the scale bar. **D)** Surface representation of PINK1-induced MPPα lid and active site engagement across HDX-MS time points, visualized in PyMOL. **E)** MPPα mutations from ClinVar within the dynamic MPPα regions identified from our HDX-MS dataset were visualized in PyMOL.

Notably, two of the most highly protected regions of MPPα were located at the top face of MPPα, which we have coined as the MPPα “lid” or “gate”. In this region, we observed protection of the glycine-rich loop (GRL), which was previously reported as a feature of yeast MPP that regulates substrate recognition (Dvořáková-Holá *et al*., 2010; Nagao *et al*., 2000; Shimokata *et al*., 1998). Within this lid region, we observed the most significant protection within a novel loop-helix-loop immediately after the GRL, which we have coined the RLY-helix loop, named according to the peptides in our HDX-MS dataset. This protected RLY-helix loop peptide also extended into a portion of one of the β-strands that lie just below the MPPα lid, forming a β-sheet that constitutes the upper part of the MPPα substrate binding site. The rest of this β-strand and the following strand displayed significant protection, albeit to a lesser degree. Closer inspection of the MPPα lid region reveal two other regions which re-arrange upon PINK1 binding – namely, portions of the PRQ-helix, which lies next to the RLY helix, and the short HIQT-helix, which lies next to the PRQ-helix **(Fig 4B)**. On the opposite face of MPPα, helices which form the distal part of the MPPα active site were also protected in the presence of PINK1_1-45_ – namely the NLE helix-loop-helix and RSR loop-helix **(Fig 4A, Fig S5B)**. Combined with our observations of the lid and active site β-sheets, these findings suggest that PINK1_1-45_ induces extensive conformational re-arrangements within the MPPα lid, but also binds within the MPPα substrate binding site.

To investigate the temporal progression of MPPα conformational changes following PINK1 binding, we analyzed the deuterium incorporation of specific MPPα features across the labelling timepoints. We stratified the conformational changes into two distinct stages, according to regions of MPPα which were significantly protected within 10 secs, representing the earliest stage of substrate binding, and those which were protected within 1 min to 5 min, which we posit to represent subsequent active site engagement by PINK1. In the first stage, we observed rapid protection of distinct components of the MPPα lid upon D_2_O exposure, namely the RLY helix-loop, GRL, and to a lesser degree the LMG loop **(Fig 4C)**. These observations suggest that the earliest stage in MPPα substrate recognition likely involves a large conformational shift within the GRL and RLY helix-loop to accommodate an incoming substrate. Notably, while our AlphaFold model confidently predicts the RLY helix (a.a. 362-370), this region displayed higher D_2_O exchange rates in the absence of substrate when compared to other solvent accessible helices within MPPα **(Fig S5A)**. This enhanced deuterium incorporation suggests that the lid region of MPPα exists in a highly dynamic state prior to substrate engagement, poised to re-arrange for substrate capture.

In the second stage, we observed significant protection within 1-5 mins for a subset of MPPα features which comprise the active site, namely the NLE-hinge, the RSR-helix-loop, and portions of the two active site β-strands below the lid, in addition to the sites in the first stage **(Fig 4C, Fig S5B)**. These conformational changes suggest that PINK1 also binds within the MPPα active site, and that these changes may be secondary to the lid opening. In terms of MPPβ conformation, the majority of MPPβ was unchanged in the presence of PINK1, except for one EIE-β-strand (a.a. 122-130) which lies next to the coordinated Zn^2+^ atom and parallel across the active site to the first RSY-strand in MPPα **(Fig S6B)**. This MPPβ feature displayed only a minor decrease in deuterium uptake (> - 4%) across all timepoints. Taken together, we propose a model in which MPPα-substrate binding proceeds in two putative stages: first, lid opening leading to substrate capture, and second, active site engagement **(Fig 4D)**. Further studies will be needed to assess whether protection of this region is secondary due to its proximity to the MPPα lid opening, or whether it is truly required for early substrate binding.

### Structural modelling of MPP-PINK1 and MPPα disease mutations

To further rationalize our HDX-MS results and attempt to explore how PINK1_1-45_ might trap MPP dimers, we generated AlphaFold3 models of MPPαβ in complex with either PINK1_1-45_ or MDH_1-24_ and mapped the HDX-MS % incorporation at 5 min on these structures to identify plausible binding modes **(Fig S7A)**. Despite low per-residue confidence scores for the MTS peptides, MTS binding poses converged inside the MPPαβ catalytic pocket **(Fig S7A)**, with PINK1_1-45_ interacting extensively with MPPα proximal to its NLE-hinge and substrate binding site β-strand. Strikingly, PINK1 R27 (the R-2 cleavage motif) is misplaced far from the catalytic Zn^2+^ within the MPPβ active site, which might partially explain the slow MPP turnover of PINK1. By contrast, AF3 predicts MDH_1-24_ in an extended conformation, with most of its contacts on the MPPβ side and its R-2 motif positioned in a cleavage competent position **(Fig S7A)**, similar to that observed in the crystal structure of the MDH N-MTS bound to the yeast MPP (Taylor *et al*., 2001), As such, these models likely represent an “end state” or closed conformation of MPP-MTS binding, after the GRL/lid have recruited MTSs into the catalytic pocket for cleavage.

We next examined whether disease-associated MPPα mutations might localize to some of the dynamic structural features outlined above **(Fig 4E)**. Strikingly, most MPPα mutations clustered within the lid region of MPPα, including four mutations directly within the RLY-helix-loop (Y364C, L365F, V367M, and Y375F), one within the GRL (G356S), and four within the PRQ-helix (D397N/Y, R402G/P). In terms of the MPPα substrate binding site, two mutations were found within the active site β-sheets below the RLY-helix-loop (A377T, L389I), three within the NLE-helix loop (M436I, V444M, E447K), and two within the RSR-loop-helix (R457H, E464K). MPPα E447K is particularly intriguing given its side chain orientation towards the active site, where introduction of a positively charged residue could abrogate active site binding to positively charged MTSs. Taken together, these findings suggest that our two-stage model captures fundamental mechanisms of MPPα substrate recognition, extending to substrates beyond PINK1, as loss of PINK1 cleavage alone cannot account for the severe ataxic and neurodegenerative phenotypes caused by these mutations. Future studies focusing on the differential sensitivity of MPP substrates to disease-associated mutations across cell types will be crucial to unveil the precise mechanisms of pathogenesis in these diseases.

## Concluding remarks

Our findings establish a workflow for the purification of human MPP, provide a broadly applicable IQF-FRET assay to probe MPP activity, reveal the precise MPP cleavage site on PINK1 between Ala28 and Tyr29, and demonstrate that MPP cleavage of PINK1 is not a prerequisite for PARL processing in cells. We further characterize the MPP-PINK1 interaction, highlighting its slow turnover and ability to promote MPPαβ dimerization. We utilize PINK1 as a model substrate to study the conformational changes of the active MPP dimer during substrate recognition, revealing a novel lid re-arrangement mechanism of MPPα. Overall, our work lays the foundation for future studies of MPP-substrate interactions and sheds light on the determinants of PINK1 import.

## Author contributions

**Andrew N. Bayne:** Conceptualization; investigation; visualization; writing - original draft; writing - review and editing. **Danielle M. Simons:** Investigation; visualization; writing – review and editing. **Naoto Soya:** Investigation; visualization; writing – review and editing**. Gergely Lukacs:** Conceptualization; supervision; writing – review and editing. **Jean-François Trempe:** Conceptualization; supervision; writing - original draft; writing—review and editing.

## Acknowledgements

This work was supported by an NSERC grant (RGPIN-2022-04042) and Canada Research Chair (Tier 2) in Structural Pharmacology to J.-F.T. A.N.B. was supported by a CIHR Doctoral Fellowship and a Centre de Recherche en Biologie Structurale (CRBS) Maximilian Eivaskhani in Memoriam Graduate Studentship. Mass photometry work was conducted at the Structural Biology Platform of the Université de Montréal (RRID:SCR_022303), funded by a grant from the Canada Foundation for Innovation (#30574) and now supported by the Centre d’Innovation Biomédicale of the Université de Montréal. Intact and peptide mass spectrometry assays were performed at the McGill Pharmacology SPR-MS facility, funded by a grant from the Canada Foundation for Innovation (#229792).

## Materials and methods

### Expression and purification of human MPP from E. coli

Codon optimized sequences of MPPα (a.a. 35-525) and MPPβ_his6_ (a.a. 46-489) were gene synthesized and were cloned into various vectors (pGEX-6p1, pRSF-Duet-1, pET-Duet-1) using restriction digests or Gibson Assembly (NEB). N-terminal deletions to produce Δ60-MPPα were achieved using Gibson Assembly (NEB). His-tags were removed from MPPβ and added to MPPα using the Q5 Mutagenesis Kit (NEB). Following cloning, all plasmids were verified using whole plasmid sequencing (Plasmidsaurus). For cloning and plasmid preparations, plasmids were transformed into NEB 5-alpha competent *E. coli* and isolated using QIAprep Spin Miniprep or Maxiprep Kits. For protein expression, plasmids were freshly transformed into BL21(DE3) competent *E. coli* (NEB). BL21 cells were grown in Luria Broth at 37 °C to OD_600_ 0.6-0.8, were induced with 0.1 mM or 0.5 mM IPTG, and were incubated for 12-16 hours at 16 °C, unless otherwise indicated in the manuscript. For Δ60-MPPα_his6_β his-tagged purifications, cells were harvested and lysed via sonication in lysis buffer (50 mM HEPES-NaOH pH 7.5, 300 mM NaCl, 10 % glycerol, 0.1 mg/ml lysozyme, 25 μg/ml DNase I, 5 mM MgSO_4_, cOmplete protease inhibitor tablet EDTA-free). Lysates were clarified by centrifugation at 18,000 rpm for 45 mins. His-tagged protein capture was performed using a HiTrap TALON crude (Cytiva) column on an ÄKTA pure 25 equilibrated in buffer A (50 mM HEPES-NaOH pH 8.0, 300 mM NaCl, 5 % glycerol). Lysates were loaded onto the HiTrap column at 1 mL / min in 8% buffer B (50 mM HEPES-NaOH pH 8.0, 300 mM NaCl, 250 mM imidazole, 5 % glycerol). The columns were washed in buffer A and 8 % buffer B until baseline 280 nm absorbance and proteins were eluted across a 20 min gradient from 8% to 100% buffer B. Eluted protein was concentrated using 4 mL Amicon-Ultra concentrators (10-kDa cutoff; EMD Millipore). Eluates were resolved via gel filtration on a Superdex 200 Increase 10/300 GL (Cytiva) at 0.5 mL / min in 20 mM HEPES-NaOH pH 7.5, 100 mM NaCl, 1 mM TCEP). Designated peaks were then pooled, concentrated using Amicon Ultra 0.5 mL Centrifugal Filters (10-kDa cut-off), and frozen at -80 °C as aliquots. All lysis, purification, and chromatography steps were performed at 4 °C.

During our optimizations, we noticed colony-to-colony variability in MPPβ expression **(Fig S1C)**, which we hypothesized may be due to selective pressure induced by the high copy number pRSFDuet-1 plasmid. This variability led us to screen different co-expression vectors of varying copy numbers and to utilize the pETDuet-1 vector for the rest of the studies **(Fig S1D)**. Still, even with these modifications, the amount of soluble MPPαβ dimer after purification was only 30-50 µg per L of *E. coli* culture. These yields were not improved by the typical techniques used to increase soluble protein expression in *E. coli*, including autoinduction media or osmolyte addition (Leibly *et al*, 2012; Studier, 2005) **(Fig S1E, Fig S1F)**.

### MPP activity assays using mass spectrometry

Synthetic MTS peptides were ordered from BioBasic at a minimum of 80 % purity. Peptides were re-suspended in ddH2O, and concentrations were measured on a DeNovix DS-11 spectrophotometer, by absorbance at 280 nm (amino acid sequence permitting) or 205 nm. For 205 nm concentration calculations, the extinction coefficient of each peptide was calculated according to a previously published protocol (Anthis & Clore, 2013). For LC/MS MPP cleavage time-course assays, 0.1 µM or 0.5 µM Δ60MPPα_his6_β dimer was incubated with 10 µM synthetic peptide in reaction buffer (20 mM HEPES-KOH pH 7.5, 1 mM TCEP) and incubated at 37 °C. To stop the reaction at specific time-points, 10 µL of each reaction was incubated with 10 µL of 2X stopping buffer (1 % formic acid / 10 % acetonitrile), spun down, and kept on ice. For all other cleavage assays, MPP and substrate concentrations were varied as indicated in the manuscript, incubated at either 25 °C or room temperature, and reactions were stopped by addition of stopping buffer to a final concentration of 0.5 % formic acid / 5 % acetonitrile. Eluents were analyzed on a Bruker Impact II Q-TOF mass spectrometer equipped with an Apollo II ion funnel ESI source. Data were acquired in positive-ion profile mode, a capillary voltage of 4,500 V, dry nitrogen heated at 200°C, and a mass range of 200 – 3000 m/z. Spectra were analyzed using the software DataAnalysis (Bruker). For mass spectrometry analysis, samples were captured onto an ACQUITY UPLC BEH C18 Column, 130Å, 1.7 µm, 3 mm X 100 mm column in 0.1 % formic acid / 5 % acetonitrile at a flow rate of 40 µL / min. Peptides were eluted with a 15 % to 40 % acetonitrile gradient over 25 min, followed by a washing step at 90 % acetonitrile for 6 mins, and a re-equilibration step at 5 % acetonitrile for 14 mins. Eluates were analyzed on a Bruker Impact II Q-TOF mass spectrometer equipped with an Apollo II ion funnel ESI source. Data were acquired in positive-ion profile mode, a capillary voltage of 4,500 V, dry nitrogen heated at 180°C, and a mass range of 150 – 2200 m/z. Spectra were analyzed using the software DataAnalysis (Bruker). Total ion chromatograms were manually inspected for expected full-length peptides and cleavage products. Extracted ion chromatograms (EIC) were generated for all reactant and product peptides with a mass tolerance of 0.01 Da and EIC intensities in each run were exported for further data processing and visualization in GraphPad Prism.

### MPP activity assays using internally quenched MDH-like peptides

Internally quenched fluorescent peptides were ordered from BioBasic at a purity of > 95%. For Km determination, P4K peptide (MLSALARPASAAK(Abz)RRSFSTSAQ(Dnp)) in varying concentrations was incubated with 0.05 µM MPPαβ dimer in 20 mM HEPES-KOH pH 7.5 in 96-well Nunc™ F96 MicroWell™ Black polystyrene plates (ThermoFisher). Fluorescence was measured on a Tecan Spark multimode plate reader with an excitation wavelength of 320 nm and an emission wavelength of 420 nm. Gain settings were obtained for each plate by measuring substrate fluorescence in the absence of enzyme for 5 min, and then applying the optimal gain value for the plate to the subsequent measurement. For PINK1 peptide screens, IC_50_ determination, and divalent cation screens, 10 µM P4K peptide was incubated with 0.05 µM MPPαβ dimer in 20 mM HEPES-KOH pH 7.5 in 96-well Nunc™ F96 MicroWell™ Black polystyrene plates (ThermoFisher). For IC_50_ determination, fluorescence was measured using the Tecan Spark, as stated above, except with a manual gain value of 165. For PINK1 peptide and cation screens, fluorescence was measured on a PerkinElmer EnSpire Multimode Plate Reader with an excitation wavelength of 320 nm, an emission wavelength of 420 nm, a measurement height of 8 mm, and 100 flashes. All reactions were performed at room temperature (22°C) and were monitored with plate readings every 10 seconds. Resulting curves were exported and processing using in-house Python scripts for non-linear curve fitting and v_0_ determination. IC_50_ calculations were performed on normalized v_0_ values (% activity) in GraphPad Prism 10 using nonlinear curve fitting with variable slope.

### Cell culture and immunoblotting

U2OS PINK1 KO cell lines were a gift from the Fon and Durcan labs. Cells were cultured at 37 °C / 5% CO2 in DMEM supplemented with 10% fetal bovine serum and 1x penicillin-streptomycin. pCMV(d1)TNT PINK1(WT)-3HA, denoted as PINK1-HA throughout the manuscript, was obtained from Noriyuki Matsuda. Q5 site-directed mutagenesis (E0554) kits from New England Biolabs were used to generate the corresponding PINK1 mutants used in this study. Cells were transfected using Lipofectamine 3000 according to the manufacturer’s instructions for 24 hours. After 24 hours, cells were treated with DMSO, CCCP, or MG132 at the denoted concentrations and timepoints. Following treatment, cells were washed twice with 20 mM HEPES pH 7.5, 100 mM NaCl, harvested, pelleted at 500 g for 5 mins, and were re-suspended in lysis buffer (20 mM HEPES pH 7.5, 100 mM NaCl, 1% Triton-X-100, 0.2 % SDS, supplemented with Halt™ Protease Inhibitor Cocktail (ThermoFisher) and PhosSTOP phosphatase inhibitors (Roche). Samples were incubated on ice for 30 mins, and then were spun at 16000 g for 30 mins. Clarified lysates were collected and protein concentrations were measured with the Pierce™ BCA Protein Assay according to manufacturer’s instructions. Samples were heated at 72 °C for 10 mins, and equal amounts were resolved by SDS-PAGE on 4–20% or 10% Mini-PROTEAN® TGX™ gels (Bio-Rad). For western blots, gels were transferred onto PVDF membranes. Following transfer, membranes were blocked using 5 % BSA in TBS-T (TBS + 0.1 % Tween). All primary antibodies were prepared in TBS-T and 3 % BSA and were incubated overnight at 4°C. Imaging was performed using HRP-conjugated secondary antibody (1:10000) and Clarity Western ECL Substrate (Bio-Rad). Primary antibodies used in this study were: anti-PINK1 (D8G3; Cell Signalling #6946), anti-TOM40 (E6Q3Z; Cell Signalling #55959), anti-TOM20 (D8T4N; Cell Signalling #42406), anti-ACTB (A2228; Sigma-Aldrich), anti-PMPCA/MPPα (Sigma; HPA021648), anti-PMPCB/MPPβ (Proteintech; 16064-1-AP), anti-Phospho-Ubiquitin Ser65, denoted as anti-pUb in the manuscript (E2J6T; Cell Signalling #62802).

### Intact protein mass spectrometry

1 µg of Δ60MPPα_his6_ monomer or Δ60MPPα_his6_β dimer in 0.1 % formic acid / 5 % acetonitrile were loaded onto a BioResolve RP mAb Polyphenyl Column, 450Å, 2.7 µm, 2.1 mm X 100 mm at 200 µL / min in 0.1 % formic acid / 5 % acetonitrile and were eluted with a 5 % to 90 % acetonitrile gradient over 20 min. Eluents were analyzed on a Bruker Impact II Q-TOF mass spectrometer equipped with an Apollo II ion funnel ESI source. Data were acquired in positive-ion profile mode, a capillary voltage of 4,500 V, dry nitrogen heated at 200°C, and a mass range of 200 – 3000 m/z. Spectra were analyzed using the software DataAnalysis (Bruker). The total ion chromatogram was used to determine where the protein eluted, and spectra were summed over the entire elution peak. The multiply charged ion species from 15,000 to 75,000 m/z were deconvoluted at 10,000 resolution using the maximum entropy (MaxEnt) and MaxEntX methods. Observed masses were searched against the corresponding protein sequences using Bruker SequenceEditor software.

### Mass photometry of MPPαβ

Mass photometry measurements were obtained using a Refeyn TwoMP mass photometer. Prior to sample analysis, a calibration curve was generated using sweet potato β-amylase (Sigma-Aldrich) in MPP purification buffer, to correlate ratiometric contrast to molecular mass. For MPP-MTS measurements, 100 nM of Δ60-MPPα_his6_ or Δ60-MPPα_his6_MPPβ in 20 mM HEPES pH 7.5, 100 mM NaCl, 1 mM TCEP were kept at room temperature prior to peptide addition. Immediately before mass photometry measurement, PINK1_1-45_ or MDH_1-24_ were diluted into the protein stocks at the denoted concentrations, and 2 µL of MPP-MTS or apo-MPP mixtures were pipetted into 18 µL of MPP purification buffer on a glass coverslip for MP measurement. Mass histograms were binned, fitted with Gaussian curves, and analyzed in DiscoverMP software (v2024 R1, Refeyn).

### Hydrogen-deuterium exchange mass spectrometry of MPPαβ

HDX-MS experiments were carried out as described (Soya *et al*, 2019). Briefly, 15 μM of Δ60-MPPα_his6_MPPβ in 20 mM HEPES pH 7.5, 100 mM NaCl, 1 mM TCEP pre-incubated alone, or with PINK1_1-45_ peptide in 6-fold molar excess for 1 minute on ice. HDX was initiated by diluting the pre-incubated Δ60-MPPα_his6_MPPβ (apo) solution or Δ60-MPPα_his6_MPPβ + PINK1_1-45_ solution using 1:14 dilution ratio into D_2_O-based buffer (same components as H_2_O-based MPP stock solution dissolved in D_2_O. For Δ60-MPPα_his6_MPPβ + PINK1_1-45_, D_2_O-based buffer also contains 6 μM of PINK1_1-45_). The HDX incubation period and temperature were set to 10s, 1 min, and 5 min, and 22°C, respectively. HDX was quenched with chilled quenching buffer (300 mM glycine, 8 M urea in H_2_O, pH 2.4) using 3:7 dilution ratio. Quenched solutions were flash frozen in methanol-containing dry ice, and samples were stored at −80°C until use. For the undeuterated control, initial dilution was made in H_2_O-based buffer. Prior to ultra-high-performance liquid chromatography (UHPLC)-MS analysis, the deuterated MPPα_his6_β ± PINK1_1-45_ was digested on an online immobilized pepsin column prepared in-house at a flow rate of 30 µl/min for 1.5 min. The resulting peptides were trapped on a C8 EXP Trap cartridge (2.7 µm HALO, 2.1 x 5 mm; Optimized Technologies, Oregon city, OR) Following desalting for 1.5 min at a flow rate of 200 µl/min, peptides were loaded onto a Hypersil GOLD C8 LC column (1 mm x 50 mm, 1.9 µm; Thermo Fisher Scientific) connected to an Agilent 1290 Infinity II UHPLC system. Peptide separation was achieved using a 5%– 40% linear gradient of acetonitrile containing 0.1% formic acid over 8 min at a 65 µl/min flow rate. To minimize back-exchange, the columns, solvent delivery lines, injector, and other accessories were placed in an ice bath. The C_8_ was directly connected to the electrospray ionization source of an LTQ Orbitrap Eclipse mass spectrometer (Thermo Fisher Scientific), and mass spectra of peptides were acquired in positive-ion mode for m/z 200–2,000. Triplicate measurements were performed for each time point. Identification of peptides was carried out in separate experiments by tandem MS (MS/MS) analysis in data-dependent acquisition mode, using collision-induced dissociation. All MS/MS spectra were analyzed using Proteome Discoverer 2.4 (Thermo Fisher Scientific). Peptide searching results were further manually inspected, and only those verifiable were used in HDX analysis. The deuteration (%) as a function of incubation time was determined using HDExaminer 3.4 (Trajan Scientific and Medical).

## Supplemental data

**Figure S1.**
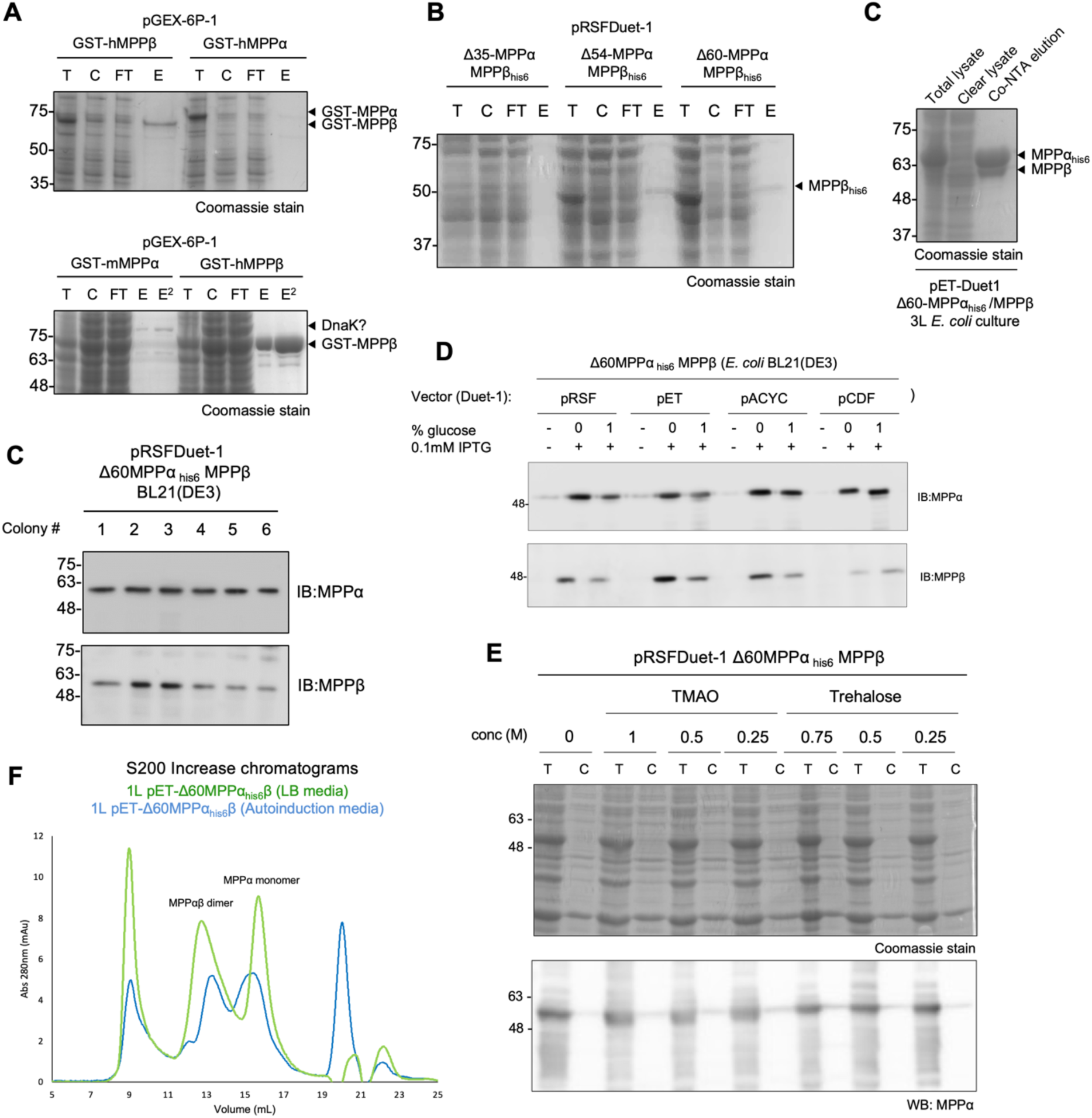
Optimization of recombinant human MPP purification from *E. coli*. **A)** MPP constructs, as denoted, were expressed in *E. coli* BL21 (DE3) cells upon addition of 0.5 mM IPTG at OD_600_ for 16 hours at 16 °C. Cells were lysed via sonication, clarified at 18000 rpm for 45 mins, and purified using Glutathione Sepharose 4B resin (Cytiva), eluted in 20 mM HEPES pH 7.5, 300 mM NaCl, 1 mM TCEP, 20 mM glutathione. Samples were mixed with Laemmli dye and resolved on SDS-PAGE. T denotes total lysate (*i.e.* soluble and insoluble), C denotes clarified lysates (*i.e.* the soluble fraction), FT denotes flow-through from affinity resin, E denotes elution, and E* denotes elution loaded at 2x concentration. **B)** Denoted constructs were expressed using 0.4 mM IPTG at OD_600_ for 16 hours at 16 °C, as above, and purified using TALON Superflow resin (Cytiva), in 20 mM HEPES pH 7.5, 300 mM NaCl, 1 mM TCEP, and 20 mM imidazole for washes and 250 mM imidazole for elution. Samples were resolved on SDS-PAGE, as above. **C)** Purification summary of Δ60-MPPα-MPPβ_his6_ from 3L of *E. coli* culture, captured using HiTrap TALON crude columns, as described in the main methods. Samples were resolved on SDS-PAGE, as above. Elutions were concentrated and resolved on gel filtration, visualized in **Fig 1B. D)** 6 colonies from the same transformation of pRSFDuet-1 Δ60MPPαβ were grown, induced, lysed, and clarified in parallel. Comparisons of soluble MPPα and MPPβ yields were measured via immunoblotting of clarified lysates. **E)** Screening of various *E. coli* co-expression vectors (pRSF, pET, pACYC, pCDF) to reduce variability in soluble MPPβ yields. *E. coli* cultures were induced via addition of 0.1 mM IPTG, either with or without 1 % glucose, which serves to reduce basal, leaky expression from T7 promoters. Soluble lysates were immunoblotted to assess yields. **F)** Trimethylamine N-oxide (TMAO) and trehalose were added to lysis buffer at varying concentrations to assess their effect on MPP solubility post-lysis. **G)** Size exclusion chromatograms comparing the yield of a 1L culture of pETDuet1-Δ60MPPαβ, in regular Luria Broth (LB media) compared to Studier’s autoinduction media.

**Figure S2.**
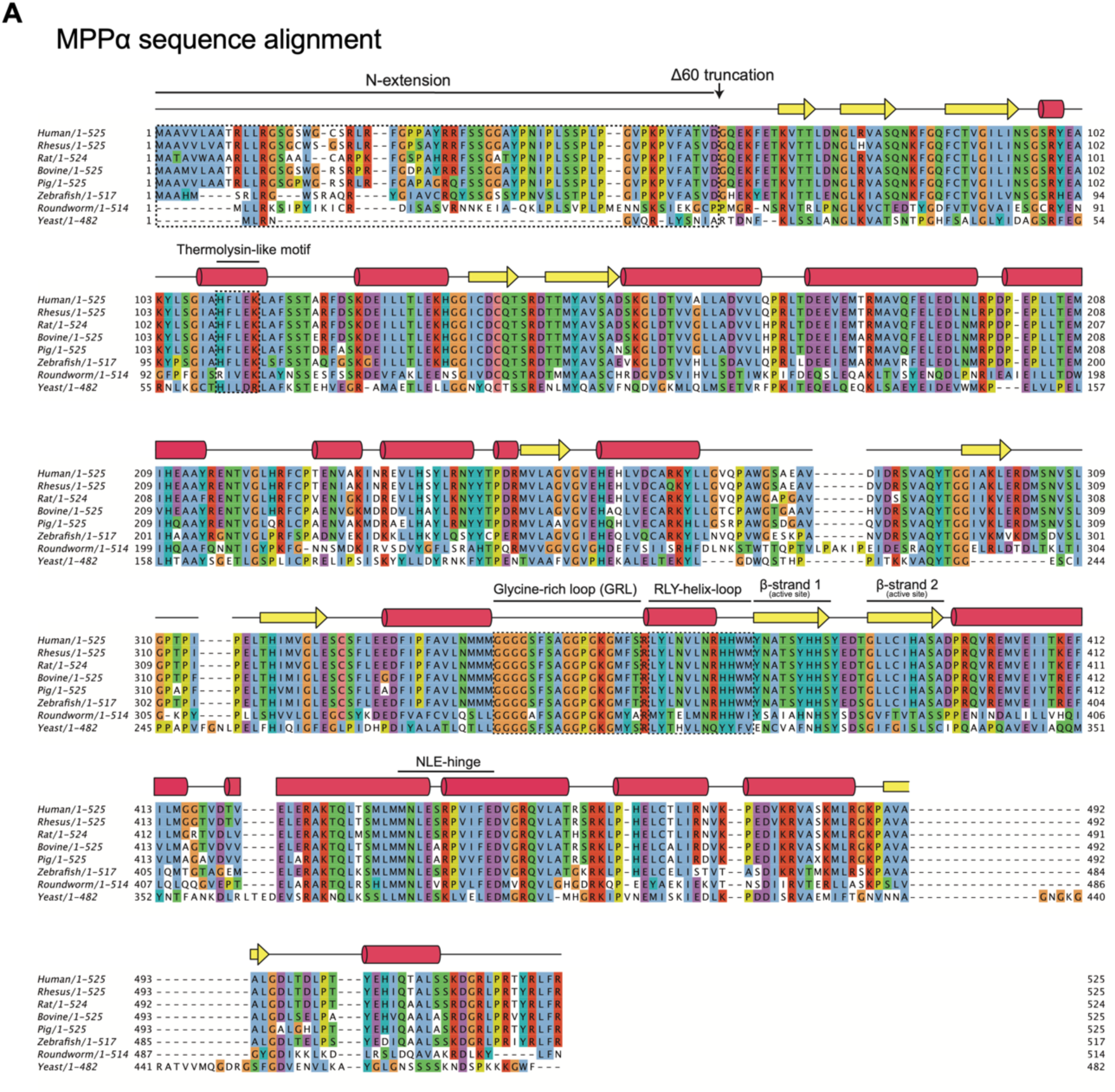
Sequence alignments of human MPPα and the PINK1 N-terminus. Selected sequences of MPPα were aligned using the MUltiple Sequence Comparison by Log-Expectation (MUSCLE) algorithm in Jalview 2.11.3.2. Alignments were annotated in Adobe Illustrator to highlight regions of interest.

**Figure S3.**
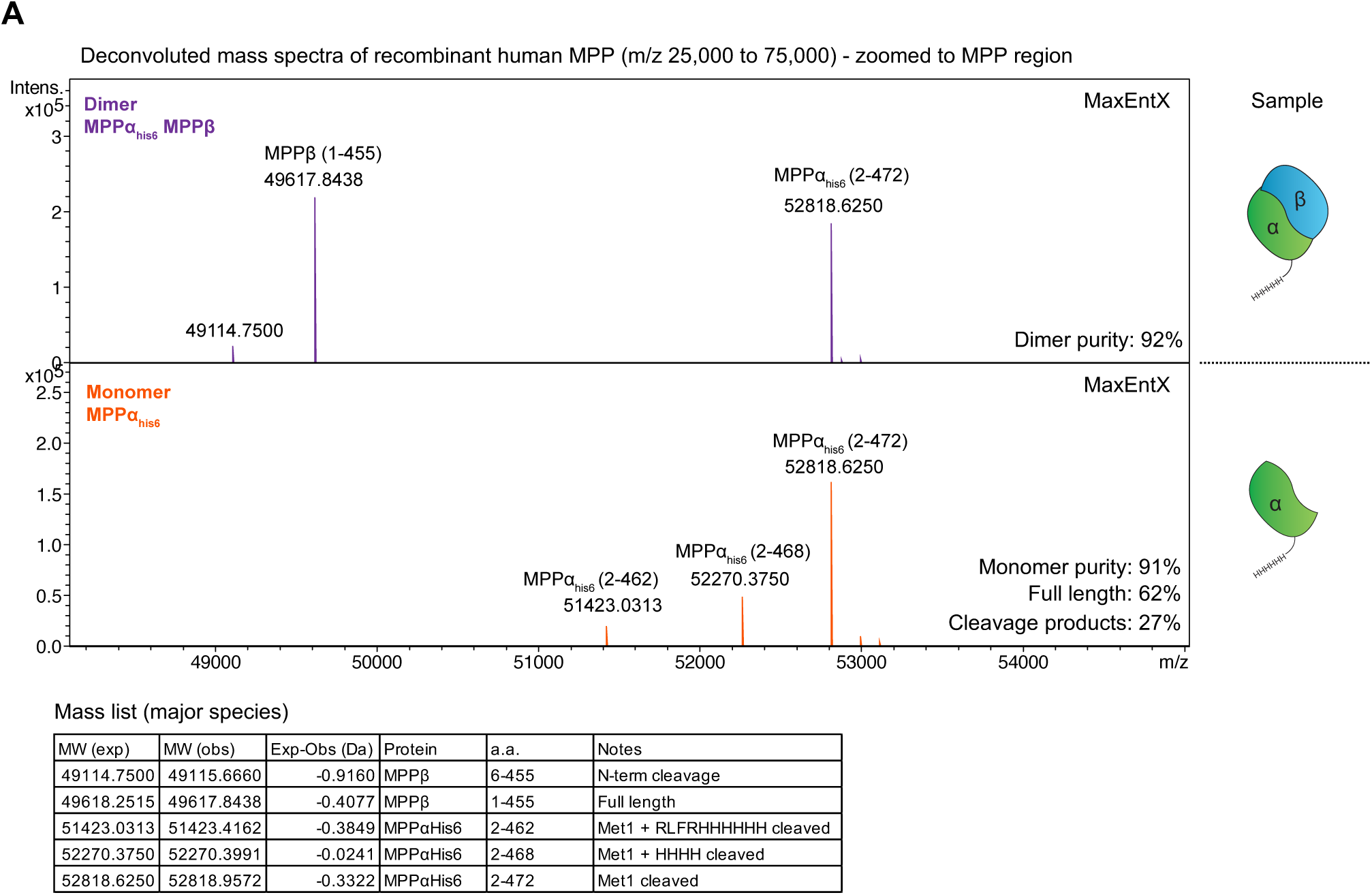
Intact mass spectrometry of purified human MPP. 1 µg of purified MPPα_his6_β dimer or MPPα_his6_ monomer were run on a Bruker Impact II Q-TOF mass spectrometer. Resulting total ion chromatograms were deconvoluted to obtain the molecular weight and were plotted as the spectra above. The observed masses for each peak were searched using Bruker Sequence Editor against the MPPαβ amino acid sequence to identify cleavage fragments (shown in the table below). Estimated purities were calculated from the intensity of a specific species divided by all the total intensity of the observed masses within each run.

**Figure S4.**
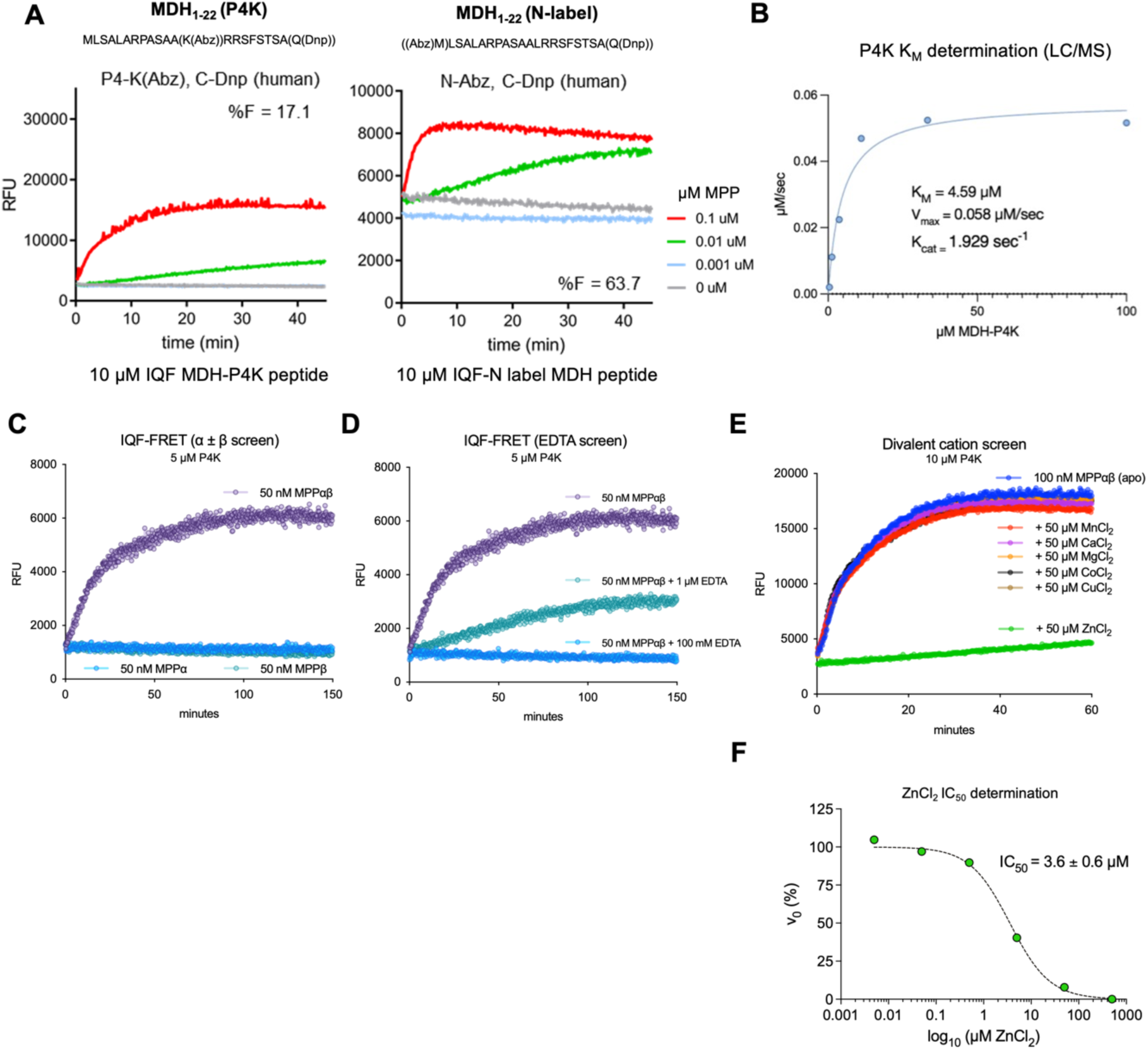
Extended characterization of MPP activity using the MDH-P4K assay and mass photometry. **A)** Comparison of P4K and N-label IQF-FRET peptides upon incubation of varying amounts of MPPαβ, as indicated. Each progress curve (n=1) was plotted in GraphPad Prism using relative fluorescence units (RFU) over time (mins). %F represents the range of fluorescence, as calculated by the initial background RFU in the absence of MPP divided by the final RFU following complete cleavage multiplied by 100. Smaller values denote higher dynamic range. **B)** P4K peptide at varying concentrations was incubated with 30 nM Δ60-MPPα_his6_MPPβ for either 2 mins at room temperature, as chosen due to the consistent linearity within the FRET progress curves, or for 90 mins at 37 °C, to generate a fully cleaved sample for standard curve generation. Following incubation, reactions were stopped by addition of 0.5 % formic acid and 5 % acetonitrile. Samples were analyzed using LC/MS on a Bruker Impact II Q-TOF mass spectrometer as denoted in the main methods. EICs were generated for MDH-P4K_1-16_ and MDH-P4K_17-24_, and a standard curve was generated using the EIC intensities of the fully cleaved 90 min samples at each P4K concentration. Intensities were converted to µM cleaved product based on this curve, and kinetic parameters were calculated using curve fitting in GraphPad Prism 10 (n=1). **C)** Monomeric Δ60-MPPα_his6,_ MPPβ, or Δ60-MPPα_his6_MPPβ were added to 5 µM of P4K peptide in 96-well, plates as described in the main methods, immediately before fluorescence measurement. Progress curves were monitored in a Tecan Spark and plotted as RFU over time. **D)** 50nM of Δ60-MPPα_his6_MPPβ was added to 5 µM P4K peptide in 96-well plates containing no EDTA, 1 µM EDTA, or 100 mM EDTA, and progress curves were immediately measured and plotted as in Fig S4C. **E)** 10mM solutions of various cations were freshly prepared in 20mM HEPES pH 7.5, before being diluted into 96-well plates containing 100 nM Δ60-MPPα_his6_MPPβ to a final concentration of 50 µM. Following 15 min incubation at room temperature, P4K peptide was added to a final concentration of 10 µM to start the reaction. Progress curves were monitored and exported as above. **F)** ZnCl_2_ was added in varying concentrations to 96-well plates, as in Fig S4E. Progress curves were exported, v_0_ were calculated from the linear portion of the reaction curve, and IC50 values were determined in GraphPad Prism 10, as described in the main methods (n=1).

**Figure S5.**
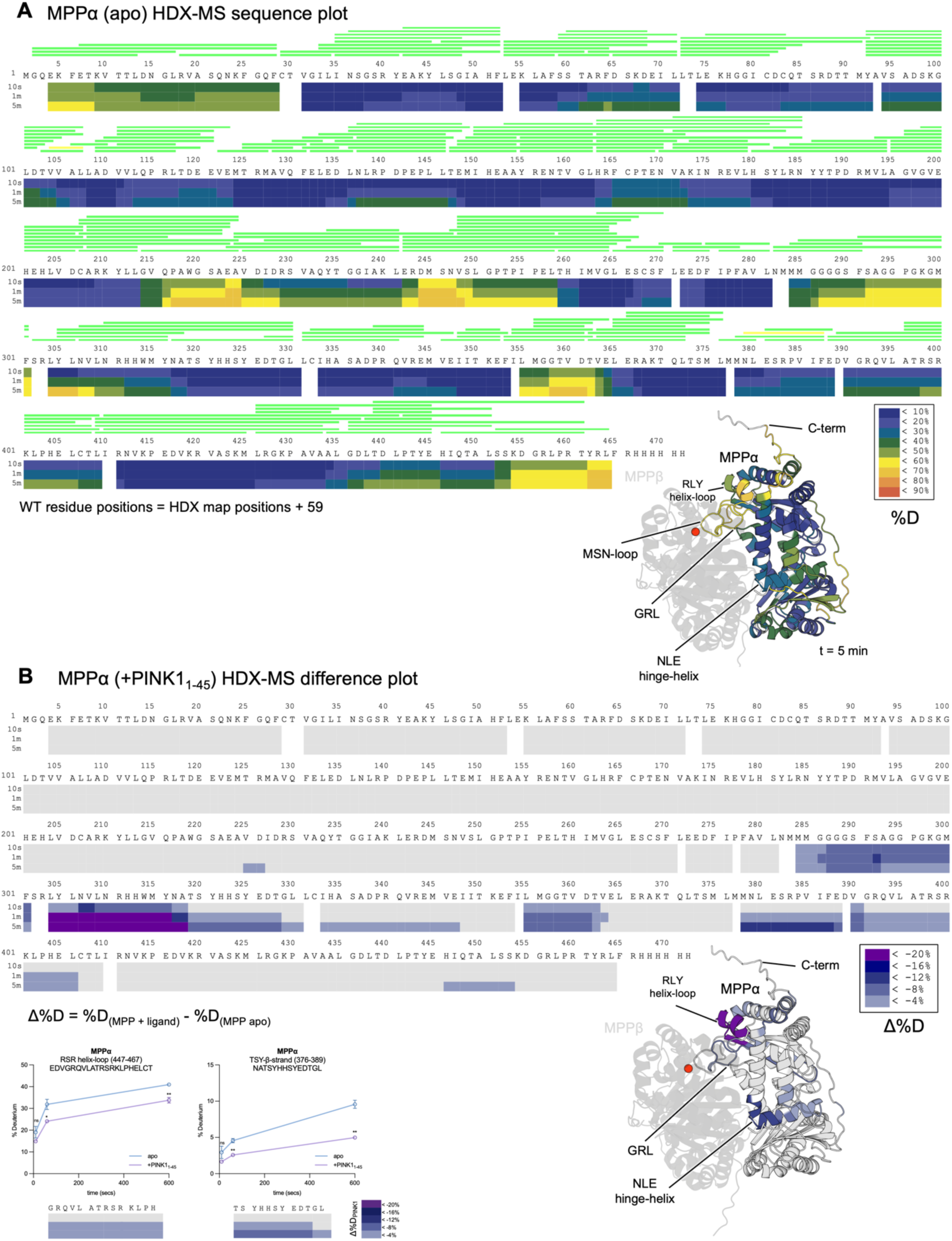
HDX-MS peptide maps of MPPα. A) Sequence map of HDX D_2_O exchange, peptide sequence coverage, and structural visualization of MPPα in the absence of PINK1_1-45_. B) Sequence map of HDX D_2_O exchange, peptide sequence coverage, and structural visualization of MPPα in the presence of PINK1_1-45_.

**Figure S6.**
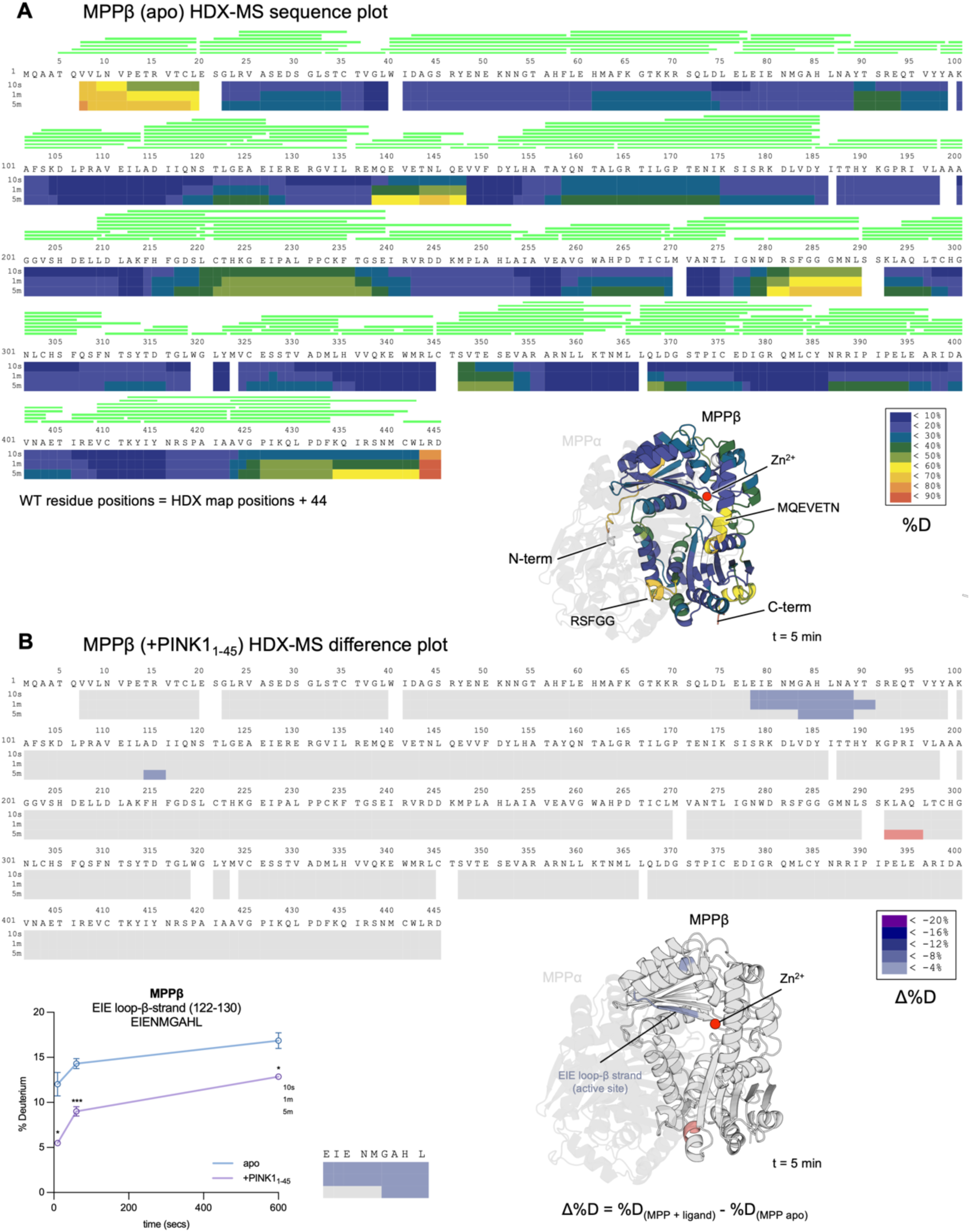
HDX-MS peptide maps of MPPβ. **A)** Sequence map of HDX D_2_O exchange, peptide sequence coverage, and structural visualization of MPPβ in the absence of PINK1_1-45_. **B)** Sequence map of HDX D_2_O exchange, peptide sequence coverage, structural visualization, and EIE-β strand time course of MPPβ in the presence of PINK1_1-45_, visualized as indicated in main methods.

**Figure S7.**
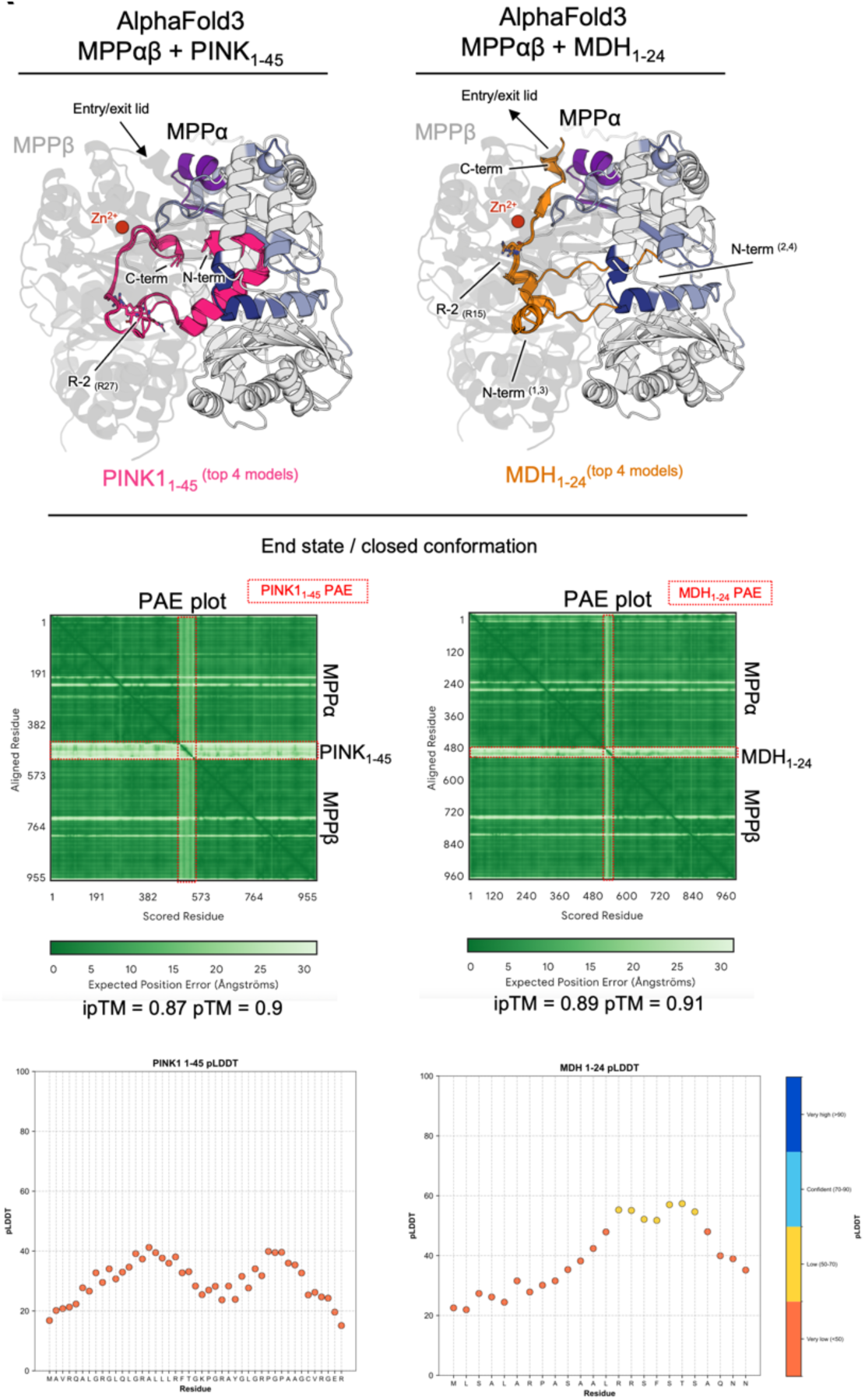
Summary of AlphaFold3 modelling of MPP-MTS complexes. Representative AlphaFold3 models of MPP-PINK1_1-45_ and MPP-MDH_1-24_, highlighting MTS topology and placement of the R-2 cleavage motif. The color scheme indicates HDX-MS protected areas (5 min) after addition of the PINK1_1-45_ peptide. PAE and pLDDT plots correspond to the highest ranked AF3 prediction.

